# Lipid membranes from naked mole-rat brain lipids are cholesterol-rich, highly phase-separated, and sensitive to amyloid-induced damage

**DOI:** 10.1101/2020.06.05.134973

**Authors:** Daniel Frankel, Matthew Davies, Bharat Bhushan, Yavuz Kulaberoglu, Paulina Urriola-Munoz, Justine Bertrand-Michel, Melissa R. Pergande, Andrew A. Smith, Swapan Preet, Thomas J. Park, Michele Vendruscolo, Kenneth Rankin, Stephanie M. Cologna, Janet R. Kumita, Nicolas Cenac, Ewan St John Smith

**Affiliations:** School of Engineering, Newcastle University, Newcastle Upon Tyne NE1 7RU, UK; Department of Pharmacology, University of Cambridge, Tennis Court Rd., Cambridge, CB2 1PD, UK; MetaToulLipidomics Facility, INSERM UMR1048, Toulouse, France; Department of Chemistry, University of Illinois at Chicago, Chicago, IL 60607, USA; Centre for Misfolding Diseases, Department of Chemistry, University of Cambridge, Lensfield Rd., Cambridge, CB2 1EW, UK; Department of Biological Sciences, University of Illinois at Chicago, Chicago, IL 60607, USA; Translational and Clinical Research Institute, Newcastle University, Paul O’Gorman Building, Framlington Place, Newcastle upon Tyne, NE2 4HH, UK; IRSD, INSERM, INRA, INP-ENVT, Toulouse University 3 Paul Sabatier, Toulouse, France

## Abstract

Naked mole-rats are extraordinarily long-lived rodents that do not develop age-related neurodegenerative diseases. Remarkably, they do not accumulate amyloid plaques, even though their brains contain high concentrations of amyloid beta peptide, even from a young age Therefore, these animals offer an opportunity to investigate mechanisms of resistance against the neurotoxicity of amyloid beta aggregation. Working in this direction, here we examine the composition, phase behaviour, and amyloid beta interactions of naked mole-rat brain lipids. Relative to mouse, naked mole-rat brain lipids are rich in cholesterol and contain sphingomyelin in lower amounts and of shorter chain lengths. Proteins associated with metabolism of ceramides, sphingomyelin and ceramide receptor activity were also found to be decreased in naked mole-rat brain lysates. Correspondingly, we find that naked mole-rat brain lipid membranes exhibit a high degree of phase separation, with the liquid ordered phase occupying up to 80% of the supported lipid bilayer. These observations are consistent with the ‘membrane pacemaker’ hypothesis of ageing, according to which long-living species have lipid membranes particularly resistant to oxidative damage. However, we found that exposure to amyloid beta disrupts the naked mole-rat brain lipid membranes while those formed from mouse brain lipids exhibit small, well-defined footprints, whereby the amyloid beta penetrates deeply into the lipid membranes. These results suggest that in naked mole-rats the lipid composition of cell membranes may offer neuroprotection through resistance to oxidative processes rather than through mechanical effects.

## Introduction

The naked mole-rat is a long-lived species having a life span that can exceed 30 years^1,2^, which is nearly ten times longer than that of rodents of comparable size. Possessing many neotenic features^3,4^ and defying many of the signs of ageing^5^, the naked mole-rat does not appear to develop neurodegenerative disease^6,7^.

Considering that amyloid beta is an age-related pathogenic agent in Alzheimer’s disease^8^, it was perhaps surprising that the naked mole-rat brain was observed to have high levels of amyloid beta, exceeding in particular those found in the commonly used 3×Tg transgenic mouse model of Alzheimer’s disease, even in the brains of young animals^6^. Somehow, therefore, naked mole-rats appear to be able to live for years with this neurotoxic peptide. Although the exact role of amyloid beta in Alzheimer’s disease (AD) remains controversial^9^ with numerous drugs targeting amyloid pathways failing in clinical trials, amyloid plaques are a hallmark of the diseased brain. It is remarkable that no such amyloid plaques are found in naked mole-rat brains despite the high levels of this peptide^6^. Thus, one would like to understand why amyloid plaques do not form in naked mole-rat brains. This question is likely to have a complex answer as in comparing the human and naked mole-rat variants of amyloid beta, one finds only a single amino acid difference, which alters the aggregation kinetics, but not the degree of toxicity to mouse hippocampal neurones^6^.

In this work we investigated whether the resistance to amyloid beta aggregation in the naked mole-rat can be traced back to specific properties of the lipid membranes of this animal, given that it is well established that lipid membranes can affect the aggregation of amyloid beta^10- 12,13^. For example, there is a strong link between cholesterol and Alzheimer’s disease, which could be related to the interaction between amyloid beta with lipid membranes^14-16^. Naked mole-rat lipids from brains and other tissue have been analysed in relation to their role in the membrane pacemaker theory of ageing, which states that long-lived species have membrane compositions that resist peroxidation, a proposed mechanism in resistance to ageing^17^. In naked mole-rat brains and within other tissues, it was observed that there was significantly less docosahexanoic acid (DHA), a fatty acid which is particularly susceptible to peroxidation, than that found in mice. In addition, it was reported that naked mole-rat membranes contain more vinyl ether linked phosphoplipids (plasmalogens) which are resistant to peroxidation.^18,19^ Plasmalogens are highly enriched in lipid rafts/microdomains and contribute to their stability.^20^ Thus, a high level of this ether-lipid may be an indicator of a high raft content, or increased raft stability.^21^

Here we used atomic force microscopy (AFM), as this method presents many advantages for examining lipid bilayer based systems, in particular its ability to quantify two dimensional ordering and to follow interactions of the membrane with peptides in real time^22,23^. More specifically, we combined AFM with lipidomics to compare naked mole-rat and mouse brain lipids in terms of composition and two dimensional ordering. We then used AFM to follow the interactions of synthetic human amyloid beta with brain-derived lipid membranes from both mouse and naked mole-rat. The membrane pacemaker theory of ageing suggests that cell membranes of long-lived species will possess more peroxidation resistant species which may also be raft forming^18^. We thus hypothesise that naked mole-rat brain derived lipid bilayers show a high degree of phase separation (two-dimensional ordering) and that there may be a unique lipid signature that protects the membranes against amyloid beta induced damage.

## Results

### Comparison of the lipid membrane compositions of naked mole-rat and mouse brains

Mass spectrometry of brain total lipid extract shows that there are key differences between naked mole-rat and mouse brains, which may have implications for amyloid beta assembly and membrane interactions. Firstly, naked mole-rat brains have significantly more cholesterol than mouse (**Figure 1A**). Also of note is the relatively low level of sphingomyelin in naked mole-rat brains relative to mouse (**Figure 1B**). Pettegrew et al. found that the levels of brain sphingomyelin positively correlate with amyloid beta plaques, however other studies show the opposite correlation between sphingomyelin and AD^24^. In contrast to the observations for cholesterol and sphingomyelin, the concentrations of the total phospholipids and ceramides were not different between naked mole-rat and mouse brains (**Figure 1C,D**). The relative abundances of these different lipid species were evaluated and presented as heatmaps (**Figure 1E,F**) and as tables (**Supplementary Tables 1-6**).

**Figure 1.**
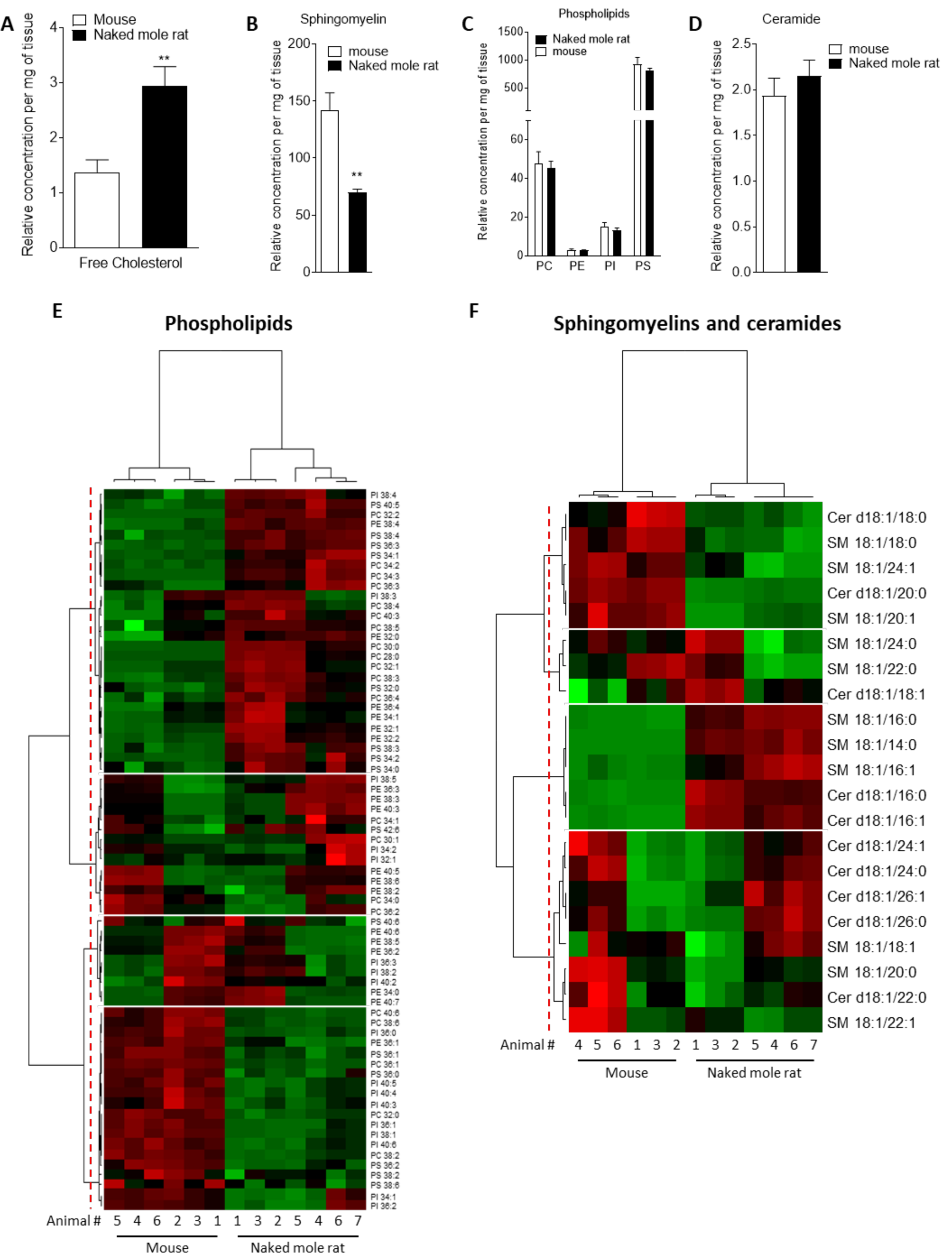
Mass Spectrometry of brain derived total lipid extract. **(A-D)** Quantification of cholesterol (A), sphingomyelin (B), phosphatidylcholine (PC), phosphatidylethanolamine (PE), phosphatidylinositol (PI) and phosphatidylserine (PS) (C), and ceramides (D) in lipid extract from mouse (white bar) or naked mole-rat (black bar) brains. Data are expressed as mean ± SEM. Statistical analysis was performed using the Mann-Whitney test. **p<0.01, significantly different from mouse group. Lipids were extracted from the brains of 7 naked mole-rats and 6 mice (2 independent experiments). **(E)** Heat map of the different phospholipids quantified by LC-MS/MS. Data are shown in a matrix format: each row represents a single phospholipid, and each column represents the lipid extract of the brain of one animal (6 mice and 7 naked mole-rat). Each colour patch represents the normalized quantity of phospholipid (row) in a single animal brain (column), with a continuum of quantity from bright green (lowest) to bright red (highest). The pattern and length of the branches in the left dendrogram reflect the relatedness of the phospholipids. The dashed red line is the dendrogram distance used to cluster the phospholipids. The pattern and length of the branches in the top dendrogram reflect the relatedness of the different animals. **(F)** Heat map of the different sphingomyelins and ceramides quantified by LC-MS/MS. Data are shown in a matrix format: each row represents a single sphingomyelin or ceramide, and each column represents the lipid extract of the brain of one animal (6 mice and 7 naked mole-rat). Each colour patch represents the normalized quantity of sphingomyelin or ceramide (row) in a single animal brain (column), with a continuum of quantity from bright green (lowest) to bright red (highest). The pattern and length of the branches in the left dendrogram reflect the relatedness of the sphingomyelins and ceramides. The dashed red line is the dendrogram distance used to cluster the phospholipids. The pattern and length of the branches in the top dendrogram reflect the relatedness of the different animals.

We found that these lipidomic profiles were different between mouse and naked mole-rat brains, as illustrated by a clustering of mouse and naked mole-rat depending on the number of carbons of the phospholipids, or of the sphingomyelins and ceramides (**Figure 1E,F**, top dendrograms). Concerning the phospholipids, four major groups of lipids were identified: Cluster 1 contained lipids which were more abundant in naked mole-rat than mouse brains, Clusters 2 and 3 contained lipids which were not differentially expressed between mouse and naked mole-rat brains, and Cluster 4 contained lipids that were more abundant in mouse brains (**Figure 1E**). However, these four clusters did not reveal a distinct pattern of species. The ceramide and sphingolipids clustering highlighted 4 clusters: Cluster 1 was composed of lipids more abundant in mouse brains, Cluster 3 by lipids more abundant in naked mole-rat brains, and Clusters 2 and 4 being no different between the two species. Interestingly, the sphingomyelins and ceramides preferentially presented in naked mole-rat brains (Cluster 3) were the shortest lipids quantified. Thus, this lipid analysis revealed differences in the lipid composition of mouse and naked mole-rat brains.

With the finding that multiple lipids were different between the two species, we sought to investigate if this was supported at the protein level. Using an untargeted differential proteomic study, we compared protein abundance between mice and naked mole-rats (Figure 2). We observed decreased levels of acid ceramidase, the enzyme responsible for ceramide degradation into sphingosine – a primary constituent of sphingolipids – and free fatty acids, in naked mole-rat cerebellar tissue (Figure 2A), and ceramide synthase 2 (catalyses the formation of ceramide with selectivity towards very long, C22:0 – C24:0 chains) was also decreased in naked mole-rats both in hippocampal and cerebellar tissue lysate (Figure 2B,C). Sphingomyelinase (sphingomyelin phosphodiesterase 3), which catalyses the hydrolysis of sphingomyelin to ceramide and phosphocholine, is also decreased in naked mole-rat hippocampal tissue (Figure 2D) and the sphingosine-1-phosphate receptor 1 is also decreased in naked mole-rat cerebellum and cerebral cortex (Figure 2E,F). The decreased expression of these proteins establishes a connection for the decreased sphingomyelin levels in naked mole-rats and also the potential atlerations of chain-length specific ceramides.

**Figure 2.**
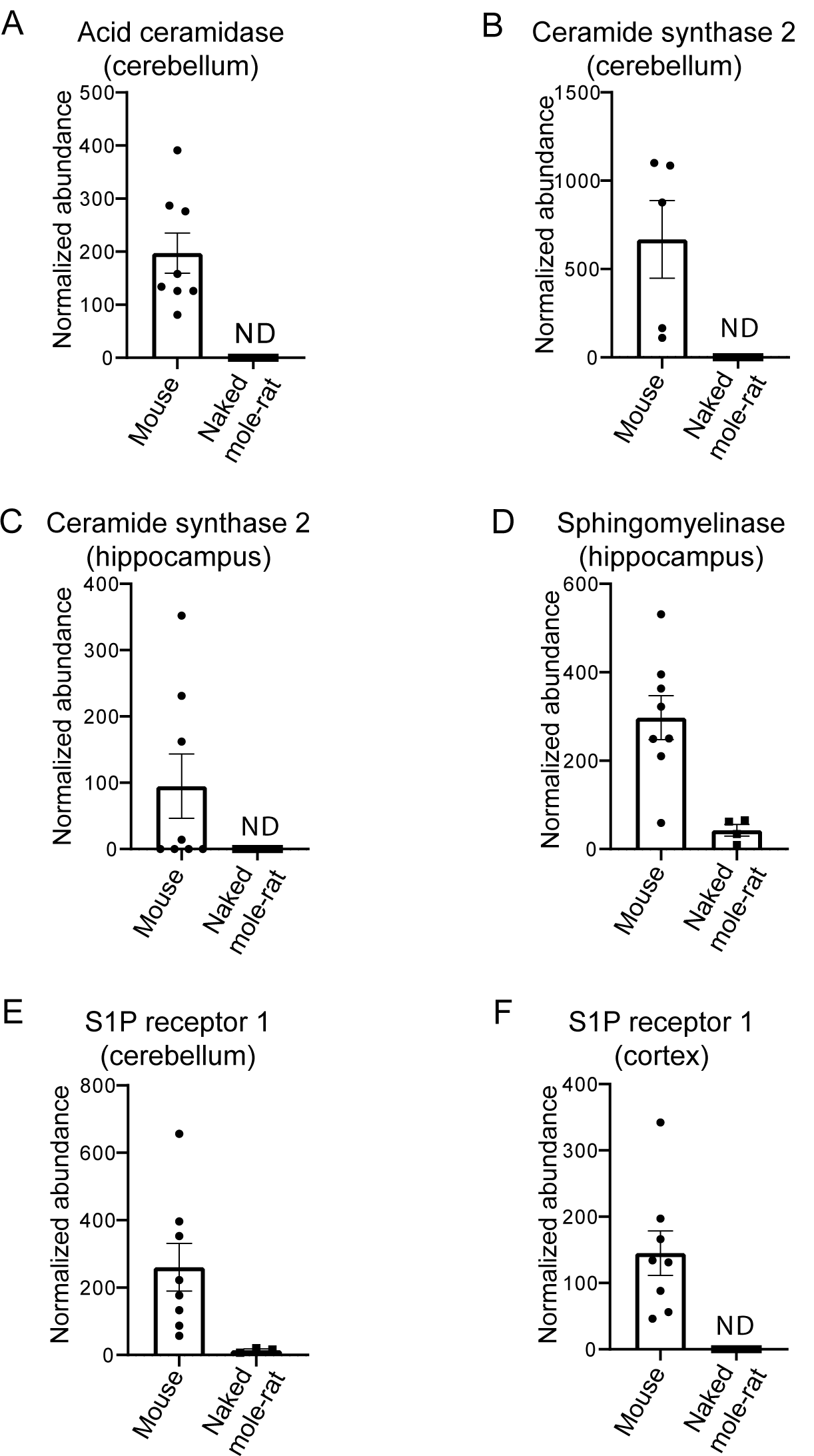
Differential proteins in mouse compared to naked mole-rats. (A) Differential levels of acid ceramidase in cerebellar tissue. Ceramide synthase 2 is decreased in naked mole-rat tissues in both the cerebellum (B) and hippocampus (C). Sphingosine phosphodiesterase 3, a member of the sphingomyelinase family is decreased in naked mole-rat hippocampus (D) where sphingosine-1-phosphate receptor 1 is decreased in the cerebellum (E) and the cerebral cortex. ND indicates not detected or quantifiable. The homology between the two species, relative to mouse, was 81% (A), 90% (B), 91.3% (C) and 95% (D).

Overall, naked mole-rat brain lipids contained a higher concentration of cholesterol and lower concentration of sphingomyelins, which were of shorter length than those present in mouse brains, but contained a higher percentage of saturated sphingomyelin (**Supp. Figure 1**). We completed the comparison of lipid composition of naked mole-rat and mouse brains by quantifying the concentration in total fatty acids (FA). No significant differences were observed between the concentrations of saturated (SFA), monounsaturated (MUFA) and polyunsaturated fatty acids (PUFA) in mouse and naked mole-rat brains (**Figure 3A-C**). However, the n-6/n-3 ratio of PUFAs was increased in naked mole-rat brains compared to mouse brains (**Figure 3D**). The relative abundances of these FA were evaluated and presented as a heatmap (**Figure 3E**) and as table (**Supplementary Table 7**). We found that the FA profile was different between mouse and naked mole-rat brains, illustrated by a clustering of mouse and naked mole-rat depending on the number of carbons within the FA (**Figure 3E**). Three major groups of FA were identified: Cluster 1 contained lipids which were more abundant in mouse brains, Cluster 2 contained FA which were not differentially expressed between mouse and naked mole-rat brains, and Cluster 3 contained lipids that were more abundant in naked mole-rat brains (**Figure 3E**). As observed for the ceramides and sphingomyelin, the SFA and MUFA abundant in naked mole-rat brains (Cluster 3) were of shorter length than those present in mouse brains. **Supplementary Tables 1 -7** provide a detailed comparative overview of the percentage of different lipid species identified in mouse and naked mole-rat brain lipids.

**Figure 3.**
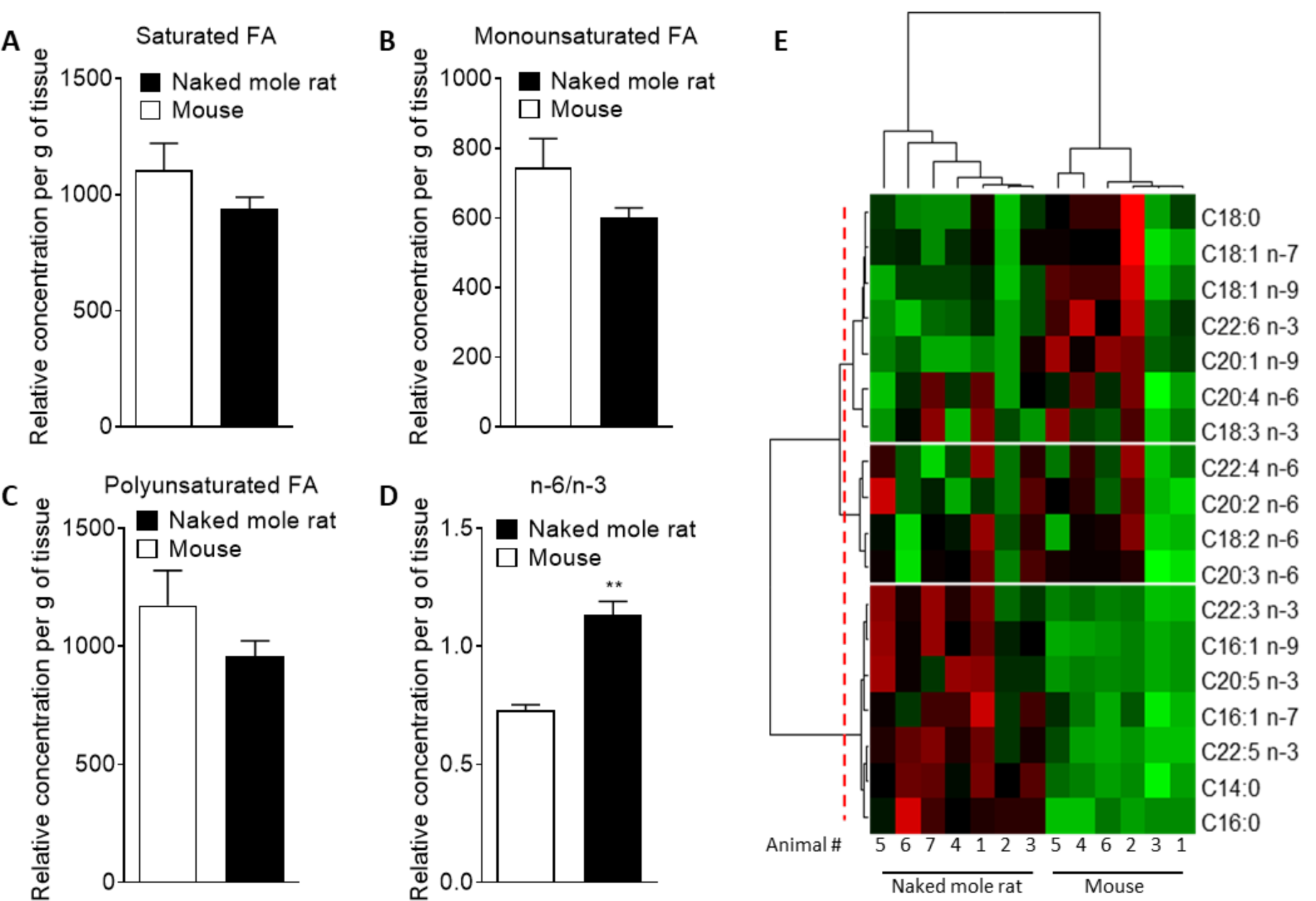
Mass Spectrometry of brain derived total fatty acids. Lipids were taken from the brains of 7 naked mole-rats and 6 mice (2 independent experiments). Quantification of (A) saturated fatty acids, (B) monounsaturated fatty acids, (C) polyunsaturated fatty acids (PUFA), and (D) n-6 PUFA/n-3 PUFA ratio extract from mouse (white bar) or naked mole-rat (black bar) brains. Data are expressed as mean ± SEM. Statistical analysis was performed using Mann-Whitney test. **p<0.01, significantly different from mouse group. (E) Heat map of the different fatty acids (FA) quantified by LC-MS/MS. Data are shown in a matrix format: each row represents a single FA, and each column represents the lipid extract of the brain of one animal (6 mice and 7 naked mole-rat). Each colour patch represents the normalized quantity of FA (row) in a single animal brain (column), with a continuum of quantity from bright green (lowest) to bright red (highest). The pattern and length of the branches in the left dendrogram reflect the relatedness of the FA. The dashed red line is the dendrogram distance used to cluster the FA. The pattern and length of the branches in the top dendrogram reflect the relatedness of the different animals.

### Amyloid beta avoids gel phase domains in model lipid membranes

Prior to an AFM examination of the interactions between human amyloid beta and the brain tissue derived lipids, we used a simple model system comprising of just two lipids, DOPC and DPPC. This system exhibits phase separation and provides a well-defined comparison for the more complex mixture of lipids present in brain derived lipids. When the binary system was mixed in a molar ratio of 3:1 of DOPC-to-DPPC, we observed the classic phase separated DPPC gel domains surrounded by the liquid disordered DOPC (**Figure 4A**). The origin of the islands of DPPC is due to the packing arrangement of DPPC chains, resulting in a height difference of around 1.5 nm to the surrounding DOPC (**Figure 4B**). Such spontaneous phase separation behaviour in model systems has been used as a rationale for the existence of lipid rafts in cells^25^. After 2 hours exposure to 8 μM of human amyloid beta the bilayer remained completely intact, but with clearly visible features in between the DPPC gel phase domains, that are absorbed on top of the DOPC liquid disordered phase (highlighted by blue circle in **Figure 4C**). These features are the amyloid beta peptide and it is to be noted that no peptide adsorbs on the gel phase domains. Upon closer inspection, the amyloid beta is observed to avoid the gel phase domains, such that the central domain is surrounded, but not covered, by the peptide (**Figure 4D**).

**Figure 4.**
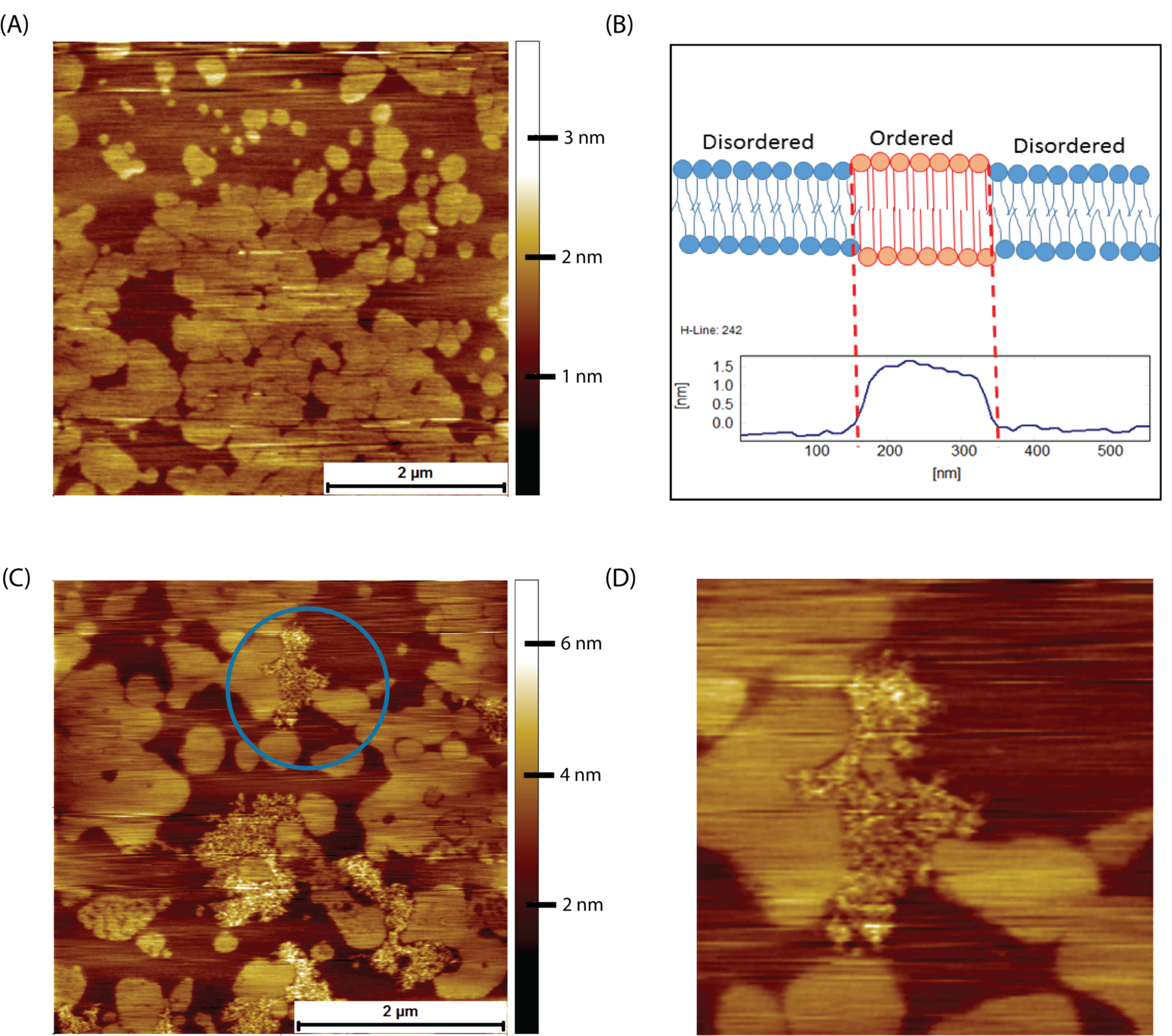
AFM contact mode images in PBS of DOPC/DPPC model supported lipid bilayers and their interactions with human amyloid beta peptide. **(A)** Supported bilayer made from DOPC to DPPC in a molar ratio of 3:1. Typical phase separation is observed with the DPPC gel phase islands surrounded by the liquid disordered DOPC phase. **(B)** Proposed origin of height difference of the gel phase relative to liquid disordered phase. **(C)** DOPC/DPPC bilayer after exposure for 2 hours to 8 μM of human amyloid beta. The amyloid beta has adsorbed exclusively onto the DOPC liquid disordered phase, the DPPC gel phase remaining untouched. **(D)** A zoom in of the area enclosed in the circle in (C)., the amyloid beta being seen to avoid the DPPC gel domains, even the small central gel domain remaining untouched.

### Lipid bilayers formed from naked mole-rat brain-derived total lipid extracts exhibit a high degree of phase separation

Lipid bilayers formed from naked mole-rat brain-derived total lipid extracts were formed on a cleaved mica surface and were found to exhibit a high degree of phase separation, with the characteristic islands (**Figure 5A**). These islands could occupy an area greater than 80% of that of the whole lipid membrane. At higher magnification, it was sometimes possible to distinguish a two-tier raft (**Figure 5B,C**), but this was not seen in all preparations. However, the high degree of phase separation was observed in lipids extracted from all naked mole-rat brains studied. For comparison mouse brain lipids exhibited little to no phase separation, such that a lipid membrane formed with little to no deviation in height and structure (**Figure 5D**).

**Figure 5.**
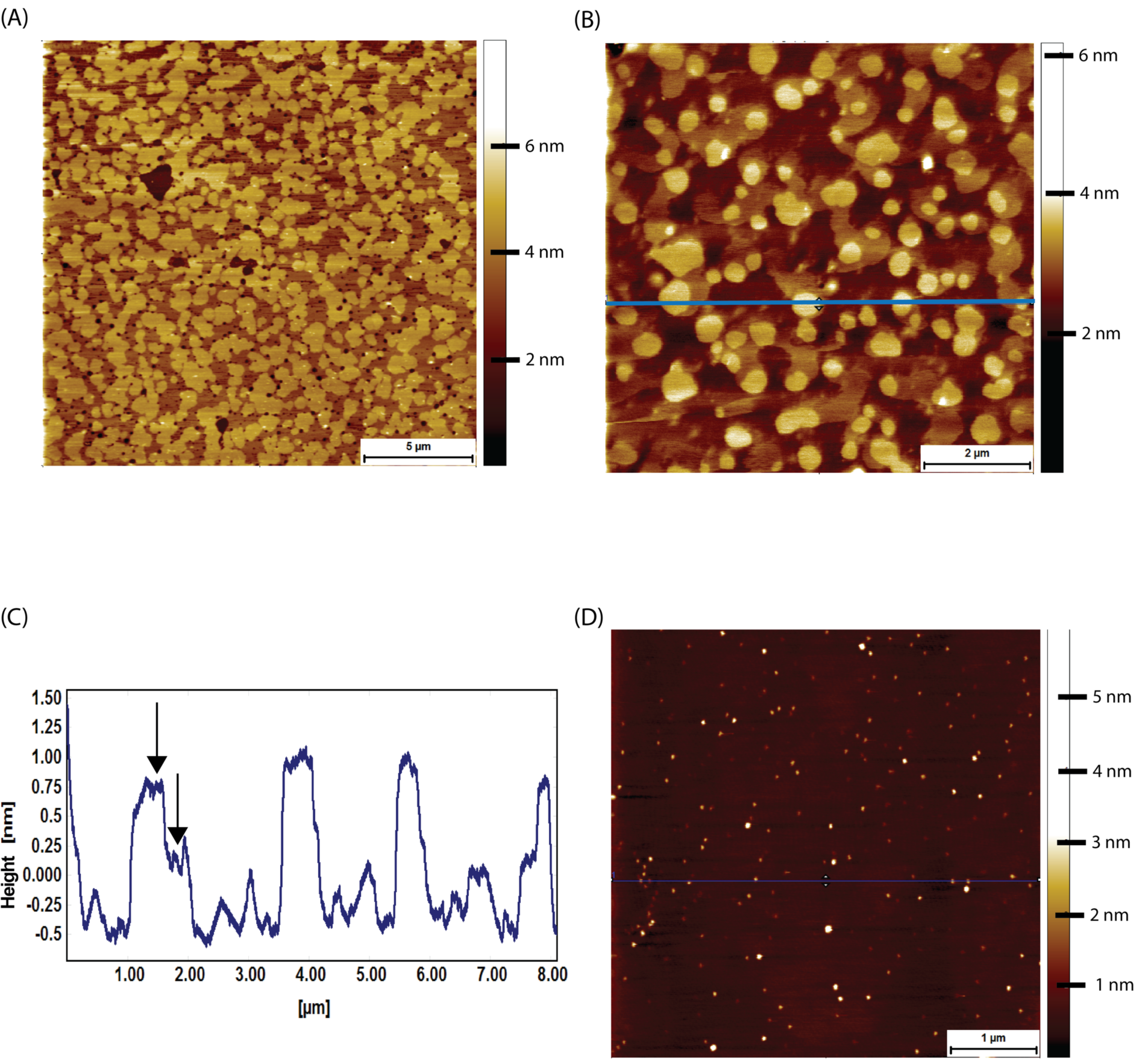
AFM tapping mode images of brain-derived supported lipid bilayers. **(A)** Supported bilayer made from naked mole-rat brain-derived lipids in PBS. More than 80% of the membrane is made up of phase separated islands of ∼1.5 nm in height relative to the surrounding liquid disordered phase. **(B,C)** Supported bilayer made of naked mole-rat brain lipids showing a two-tier raft structure, the horizontal blue line corresponding to the height profile in C, where the two-tier raft structure is evident from the peaks having a shoulder, the first at ∼0.75 nm, the second at ∼1.25 nm, indicated by arrows in the figure. **(D)** Supported lipid bilayer made from mouse brain derived lipids. In contrast to the naked mole-rat bilayers there is little evidence of phase separation.

By indenting the conical AFM tip into the lipid bilayer it is possible to measure the breakthrough force^26^, which is the force required to push through the soft thin film (**Figure 6A,B**), thereby providing an indication of the membrane fluidity. It also allows the measurement of membrane thickness, which is a necessary confirmation that a thin soft bilayer is being imaged. Although no difference was observed in the thickness of lipid bilayer membranes formed from mouse or naked mole-rat brain derived lipids (**Figure 6C**), the breakthrough force measurements for naked mole-rat membranes were significantly higher than mouse (**Figure 6D**), meaning that naked mole-rat lipid bilayer membranes are stiffer. The enhanced stiffness of naked mole-rat lipid bilayer membranes is most likely due to the increased cholesterol, which is known to increase membrane rigidity.^27^

**Figure 6.**
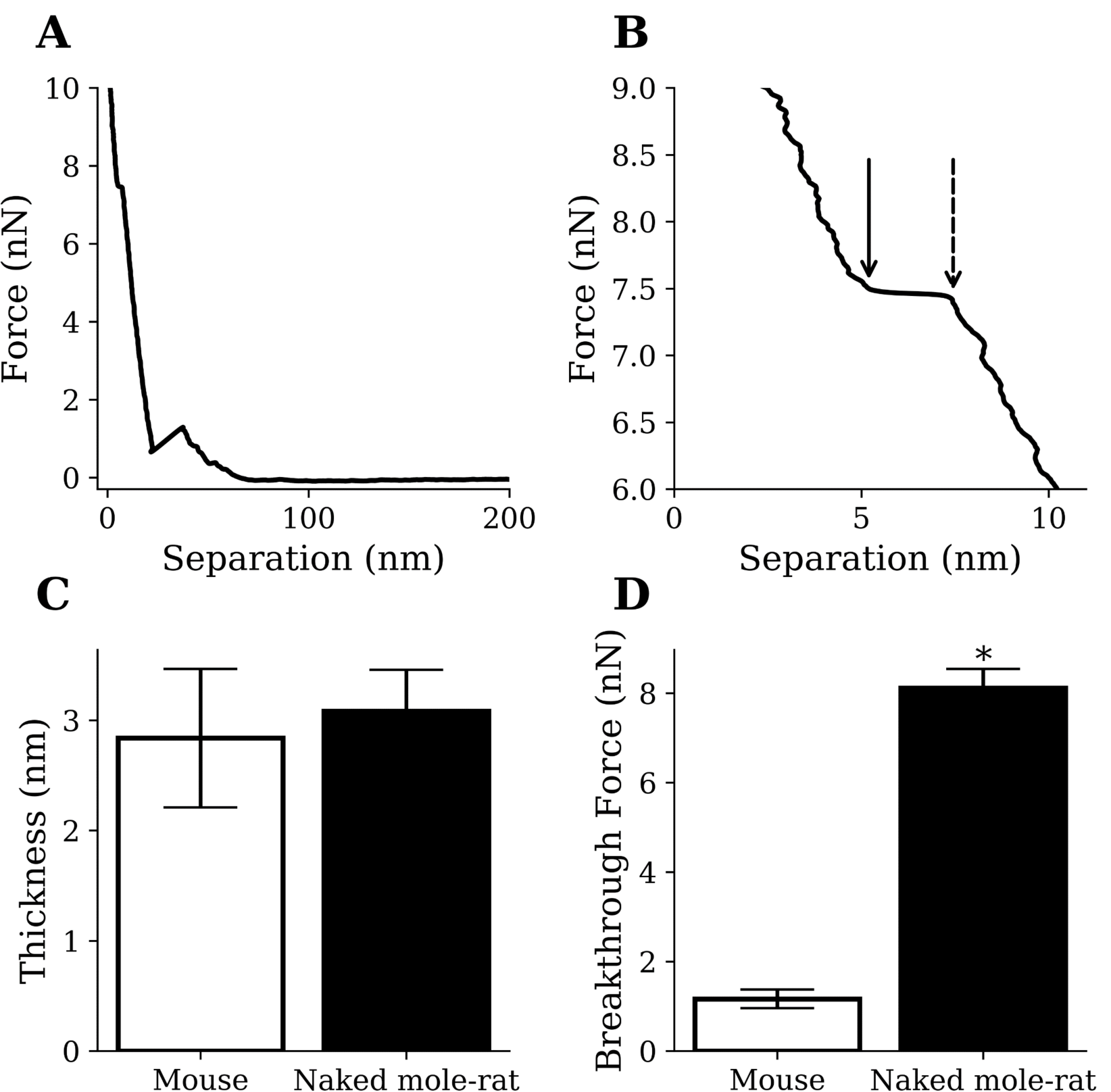
AFM results from mouse samples (n=1220) and naked mole-rat samples (n=2859) lipid bilayer indentations. **(A,B)** Example indentation on a naked mole-rat bilayer (A), where multiple breakthrough events were detected only the event closest to the mica (B) was recorded. The points indicated by the dashed and solid arrow in panel B show the start and end of the breakthrough event respectively. Thickness was recorded as the difference in separation between the points and breakthrough force was recorded at the start of the breakthrough indicated by the dashed arrow. **(C)** Thickness values of 3.1 ± 0.4 nm and 2.8 ± 0.6 nm were recorded for the naked mole-rat and mouse lipid bilayers respectively. **(D)** Breakthrough force values of 8.1 ± 0.4 nN and 1.2 ± 0.2 nN were recorded for the naked mole-rat and mouse lipid bilayers respectively. Data are expressed as mean ± SEM. Statistical analysis was performed using the unpaired t test. *p<0.05, significantly different from mouse group.

### Amyloid beta disrupts lipid bilayers formed from naked mole-rat brain-derived total lipid extracts

Prior to exposure to amyloid beta, the mouse brain derived lipid bilayer had a characteristic morphology exhibiting almost no phase separation with only the occasional, small domain being visible (**Figure 7A**). When exposed to 8 μM human amyloid beta for 2 hours, pores appeared in the mouse derived lipid bilayer (**Figure 7B**). The pores were not circular, and there was little sign of the amyloid beta that had caused them. A likely explanation is that the amyloid beta peptides have sunk into the membrane because the height profile through the pores shows that they are less deep than the thickness of the bilayer (**Figure 7C,D**). It is important to note that the integrity of the membrane, as a whole, is maintained.

**Figure 7.**
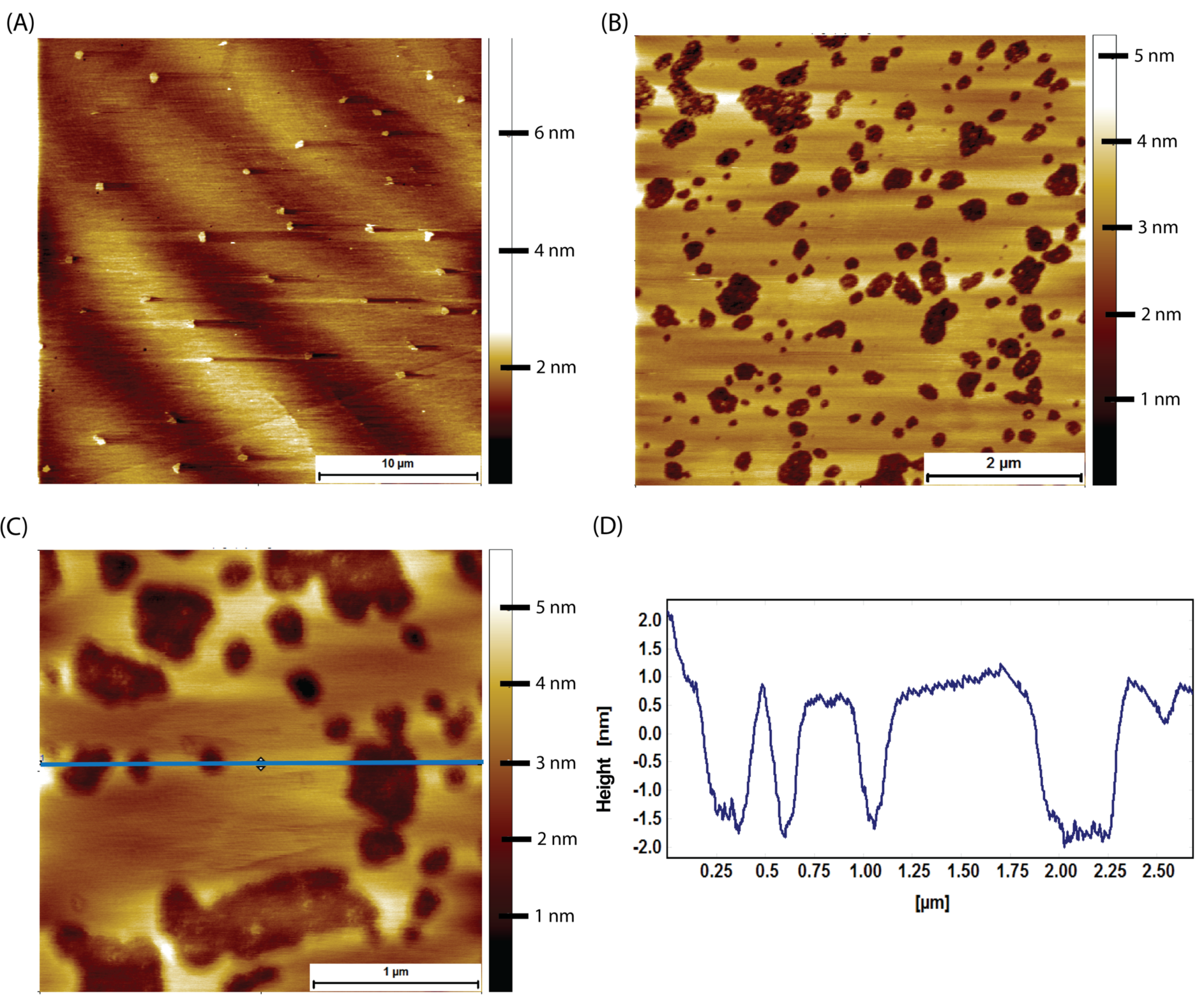
AFM tapping mode images of mouse brain tissue derived supported lipid bilayers in PBS exposed to 8 μM of human amyloid beta for 2 hours. **(A)** Large scale image showing a sparse amount of phase separation prior to amyloid beta adsorption. **(B)** Mouse brain derived lipid bilayer after exposure for 2 hours to 8 μM of human amyloid beta. Pits appear in the membrane, with some structure visible within the pits. **(C)** A close up of the pits showing that they do not break through the entire bilayer, i.e. they are not holes. **(D)** Line profile corresponding to the horizontal blue line in (C). The pits are between 2 and 2.5 nm deep, too shallow to be holes in the bilayer, which is ∼3.1 nm thick.

By contrast to what was observed with mouse lipid bilayers, upon exposure of naked mole-rat lipid bilayers to human amyloid beta under the same conditions (8 µM for 2 hours), the naked mole-rat lipid bilayer membrane is disrupted (**Figure 8**). In contrast to the raft covered membrane observed under control conditions (**Figure 8A**), incubation with amyloid beta results in the presence of only fragments of lipid bilayer (**Figure 8B,C**), > 5 nm in height relative to the surface (consistent with the thickness of the bilayer rather than the height of a raft, **Figure 8D**). Membrane fragments after amyloid beta exposure were observed with lipid bilayers formed from lipids extracted from all 3 brains examined.

**Figure 8.**
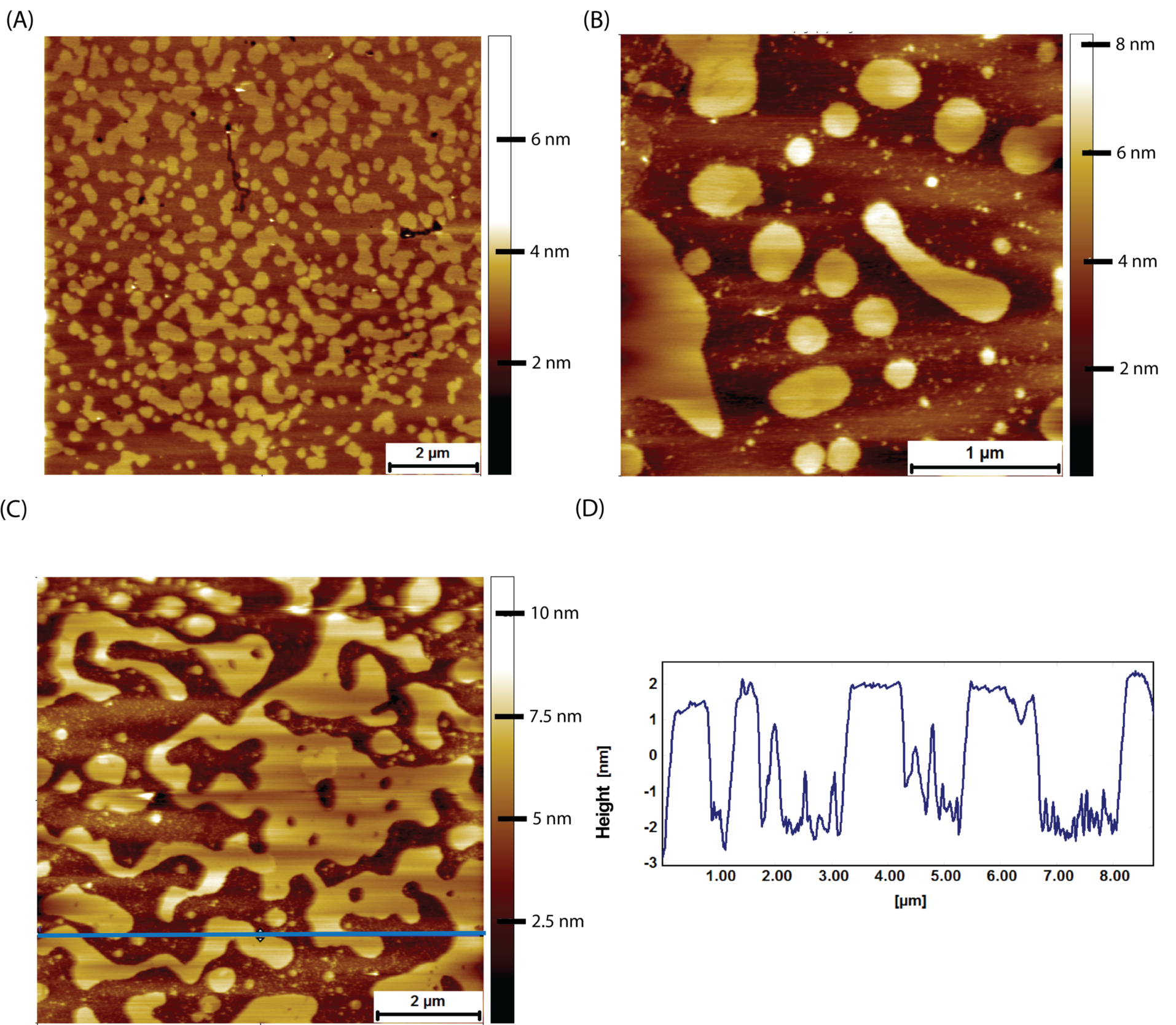
AFM tapping mode images of naked mole-rat brain derived lipid bilayers exposed to 8 μM of human amyloid beta for 2 hours. **(A)** The naked mole-rat brain derived bilayer before amyloid beta adsorption in PBS exhibiting the high degree of phase separation characteristic of naked mole-rat bilayers. **(B,C)** Naked mole-rat brain derived bilayer exposed to 8 μM of human amyloid beta for 3 hours. **(D)** A height profile corresponding to the blue horizontal line in (C), confirming the height of fragments to be of bilayer thickness of 4∽nm.

## Discussion

Our lipidomic data show fatty acids and phospholipid trends between mouse and naked mole-rat that are consistent with the previous work of Mitchell et al^19^. However, in addition we have found that cholesterol levels in naked mole-rat brains are significantly higher than in mouse brains and that sphingomyelin levels are significantly lower and of a shorter chain lengths. These lipid findings are supported by proteomic analysis of brain lysates and critical lipid metabolism enzymes were also altered in naked mole-rats compared to mice. Using AFM, we found that naked mole-rat brain lipids have a high level of phase separation, which is consistent with predictions from the membrane pacemaker theory. Regardless of whether such domains exist in a living cell, the lipids that are able to phase separate could have a role in protection against ageing. Mechanical measurements show that the extra cholesterol results in a stiffer membrane, but surprisingly, these phase-separated and stiffer membranes offer limited protection against amyloid beta induced damage. The susceptibility of naked mole-rat membranes to amyloid beta induced damage was unexpected given the amyloid beta rich environment of the naked mole-rat brains and the lack of overt neurodegeneration in this species^6^.

Cell membranes consist of more than just phospholipids, with cytoskeleton, proteins and carbohydrates all contributing to membrane stability. However, the relative difference in response between mouse and naked mole-rat lipids bilayers could be a significant contributor to the amyloid beta response in each species. It should also be noted that the total lipid extract from brain tissue will include not just lipids from cell membranes but also from mitochondria and other organelles. Interestingly, the same susceptibility of naked mole-rat membrane lipids to damage was found when using naked mole-rat amyloid beta peptide (**Supplementary Figures 2-5**). This is despite the single amino acid difference in sequence whereby histidine is replaced by arginine in naked mole rat amyloid beta. We find that naked mole rat amyloid beta readily forms fibres and fibrils (**Supplementary Figures 2 and 3**) as is the case with human amyloid beta. Again with mouse brain lipids, the naked mole rat amyloid beta sinks into the bilayer, leaving the bilayer intact but with holes (**Supplementary material Figure 4**). Upon exposure to naked mole rat amyloid beta the naked mole rat brain derived supported bilayers are broken up into membrane fragments (**Supplementary material Figure 5**). This consistency in behaviour between human and naked mole rat amyloid beta with respect to membrane interactions strengthens the argument that brain lipid composition may be critical to the physiology of neurodegeneration in naked mole rats.

In summary, naked mole-rat brain lipids have high cholesterol and low sphingomyelin levels compared to mouse brain lipids and the bilayers formed from naked mole-rat brain lipids are unusual in terms of a high degree of phase separation. By contrast, both mouse from our data and pig brain lipids^28^ show no or little signs of such two dimensional ordering. The comparative sensitivity of naked mole-rat lipid bilayers to amyloid beta damage suggests that the naked mole-rat has evolved neuroprotective mechanisms based more on the resistance to oxidative processes than mechanical resistance to support a healthy brain within an amyloid beta rich environment.

## Materials and Methods

### Animals

Experiments were performed on lipids and proteins extracted from C57BL6/J mice and non-breeding naked mole-rats. A mixture of males and females were used of both species, all adults for lipid studies. For proteomic studies, mice (N=8) were euthanized at 2-weeks (N=4) and 6 months (N=4) of age. Similarly, NMRs were euthanized at 2-4 months (N=4) and 3 years (N=4) of age. Cerebellum, cortex and hippocampus were collected and flash frozen, as previously described^29^. Mice were conventionally housed with nesting material and a red plastic shelter in temperature-controlled rooms at 21 °C, with a 12 h light/dark cycle and access to food and water *ad libitum*. Naked mole-rats were bred in-house and maintained in an inter-connected network of cages in a humidified (∼55 %) temperature-controlled room at 28 °C, with red lighting (08:00-16:00) and had access to food *ad libitum*. In addition, a heat cable provided extra warmth under 2-3 cages/colony. Mice were humanely killed by cervical dislocation of the neck and cessation of circulation, whereas naked mole-rats were killed by CO2 exposure followed by decapitation. Experiments were conducted under the Animals (Scientific Procedures) Act 1986 Amendment Regulations 2012 under a Project License (P7EBFC1B1) granted to E. St. J. Smith by the Home Office and approved by the University of Cambridge Animal Welfare Ethical Review Body. Experiments performed for proteomic analysis were done so in accordance with institutionally-approved protocols at the University of Illinois at Chicago (PI – T.J. Park).

### Lipid extraction

Lipid extraction was performed according to a modified chloroform/methanol (2:1, v/v) procedure^30^. Briefly, frozen brain tissue from naked mole-rat or mouse was crushed into small pieces by razor in a petri dish, then the sample was transferred into a 15 mL tube and 2 mL of chloroform/methanol (2:1, v/v) was added; chloroform (Sigma 288306-1L), methanol (Fisher 10675112). The suspension was vortexed and incubated at room temperature with agitation for 20 min. After the incubation, 400 µL of 0.9% NaCl was added to the mixture and was then vortexed and centrifuged at 500 x g for 10 min to separate the organic phase from the aqueous phase. The chloroform (lower) layer was transferred into a new 1.5 mL tube and was centrifuged at 500 x g for 10 min. Finally, the supernatant was collected without touching the pellet and was transferred into a new 1.5 mL tube. The sample was stored at -80 °C until the analysis.

### Brain neutral lipids analysis

Internal standards: glyceryl trinonadecanoate, stigmasterol, and cholesteryl heptadecanoate (Sigma) were added to 45 µL of lipid extract. Triglycerides, cholesterol, and cholesterol esters were analysed with gas-liquid chromatography on a Focus Thermo Electron system equipped with a Zebron-1 Phenomenex fused-silica capillary column (5 m, 0.25 mm i.d., 0.25 mm film thickness). Oven temperature was programmed to increase from 200 to 350 °C at 5 °C/min, and the carrier gas was hydrogen (0.5 bar). Injector and detector temperatures were 315 °C and 345 °C, respectively.

### Brain phospholipid, ceramide and sphingolipid analysis

Internal standards (Cer d18:1/15:0, 16 ng; PE 12:0/12:0, 180 ng; PC 13:0/13:0, 16 ng; SM d18:1/12:0, 16 ng; PI 16:0/17:0, 30 ng; PS 12:0/12:0, 156.25 ng; all from Avanti Polar Lipids) were added to 45 µL of lipid extract. Sample solutions were analysed using an Agilent 1290 UPLC system coupled to a G6460 triple quadripole spectrometer (Agilent Technologies). MassHunter software was used for data acquisition and analysis. A Kinetex HILIC column (Phenomenex, 50 × 4.6 mm, 2.6 μm) was used for LC separations. The column temperature was maintained at 40 °C. Mobile phase A was acetonitrile and B was 10 mM ammonium formate in water at pH 3.2. The gradient was as follows: from 10% to 30% B in 10 min, 100% B from 10 to 12 min, and then back to 10% B at 13 min for 1 min to re-equilibrate prior to the next injection. The flow rate of the mobile phase was 0.3 ml/min, and the injection volume was 5 μl. An electrospray source was employed in positive (for Cer, PE, PC, and SM analysis) or negative ion mode (for PI and PS analysis). The collision gas was nitrogen. Needle voltage was set at +4000 V. Several scan modes were used. First, to obtain the naturally different masses of different species, we analysed cell lipid extracts with a precursor ion scan at 184 m/z, 241 m/z, and 264 m/z for PC/SM, PI, and Cer, respectively. We performed a neutral loss scan at 141 and 87 m/z for PE and PS, respectively. The collision energy optimums for Cer, PE, PC, SM, PI, and PS were 25 eV, 20 eV, 30 eV, 25 eV, 45 eV, and 22 eV, respectively. The corresponding SRM transitions were used to quantify different phospholipid species for each class. Two MRM acquisitions were necessary, due to important differences between phospholipid classes. Data were treated with QqQ Quantitative (vB.05.00) and Qualitative analysis software (vB.04.00).

### Brain proteomic analysis

Protein extracts for each tissue (cerebellum, cortex and hippocampus) and animal were prepared and analyzed via liquid chromatography-mass spectrometry (LC-MS) analysis. Protein extracts were obtained by lysis in RIPA buffer (50mM Tris pH 8, 150 mM NaCl, 1% Triton-X 100, 0.5% sodium deoxycholate, 0.1% sodium dodecylsulfate and 0.25% NP-40) and protein concentration determined via the BCA assay (Pierce-Thermo, San Jose, CA, USA). Peptide digests of 100μg of protein were prepared for each of the tissue lysates using the filter aided sample preparation (FASP) after spiking with recombinant green fluorescent protein (GFP) into each sample at a concentration of 100fmol per 1µg protein extract as previously described^31^. Samples were desalted via C18 reverse phase chromatography and dried. All samples were resuspended in 100µL 0.1% (v/v) formic acid prior to LC-MS analysis.

### LC-MS analysis

One microliter of each sample was injected for label-free, quantitative LC-MS analysis. Mass detection was carried out with a Q-Exactive mass spectrometer (Thermo Fisher Scientific, Bremen, Germany) equipped with an Agilent HPLC system. Chromatographic separation of peptides was accomplished using a Zorbax 300SB-C18 column (3.5µm i.d. x 150mm, particle size 5µm, pore size 100Å, Agilent Technologies, Wilminton, DE). Peptides were loaded onto a Zorbax 300SB-C18 trap cartridge at a flow rate of 2µl per minute for 10 min. After washing with 0.1% formic the peptides were eluted using a 5–60% B gradient for 180 min at a flow rate of 250 nl min ^-1^ acid (mobile phase A= 0.1% formic acid; mobile phase B =0.1% formic acid in acetonitrile). The mass spectrometer was operated in a Top 10 data-dependent mode with automatic switching between MS and MS/MS with HCD fragmentation. Source ionization parameters were as follows: spray voltage, 1.5 kV; capillary temperature, 280 °C; and s-lens level, 50.0. Full-scan MS mode (*m*/*z* 300–1700) was operated at a resolution of 70 000 with automatic gain control (AGC), target of 1 × 10^6^ ions and a maximum ion transfer (IT) of 250 milliseconds. Ions selected for MS/MS were subjected to the following parameters: resolution, 17, 500; AGC, 1 × 10^5^ ions; maximum IT, 100 milliseconds; 1.5 *m*/*z* isolation window; normalized collision energy 27.0, and dynamic exclusion, 30 seconds.

### Protein identification and label-free quantification

Raw data was searched against both the SwissProt *Mus musculus* (20,239 sequences) and an in house GFP database using the Proteome Discoverer software (v2.3, Thermo Fisher, Carlsbad, CA). Trypsin was set as the protease with two missed cleavages and searches were performed with precursor and fragment mass error tolerances set to 10 ppm and 0.02 Da, respectively, where only peptides precursors of +2, +3 and +4 were considered. Peptide variable modifications allowed during the search were: oxidation (M) and deamination (NQ), whereas carbamidomethyl (C) was set as fixed modifications. Label-free relative quantitation analysis was performed across tissue types using the average of the top 3 peptides for each protein where normalization was performed across all samples and all tissues using the intensity of the GFP internal standard. Differentially expressed proteins for NMR relative to mice for each tissue type was determined by applying a Permutation test (*p* ≤ 0.05). Raw data is available at: ftp://massive.ucsd.edu/MSV000085798/raw/.

### Brain fatty acid analysis

To measure the totalbrain molecular species derivatized with a methyl ester (FAME), internal standard, glyceryl triheptadecanoate (2 μg), was added to 40 µL of lipid extract. The lipid extract was transmethylated with 1 ml BF3 in methanol (14% solution; Sigma) and 1 ml heptane for 60 min at 80 °C and evaporated to dryness. The FAMEs were extracted with heptane/water (2:1). The organic phase was evaporated to dryness and dissolved in 50 μl ethyl acetate. A sample (1 μl) of total FAME was analysed with gas-liquid chromatography (Clarus 600 Perkin Elmer system, with Famewax RESTEK fused silica capillary columns, 30-m × 0.32-mm i.d., 0.25-μm film thickness). Oven temperature was programmed to increase from 110 °C to 220 °C at a rate of 2 °C/min, and the carrier gas was hydrogen (7.25 psi). Injector and detector temperatures were 225 °C and 245 °C, respectively.

### Atomic force microscopy

AFM experiments were performed on an Agilent 5500 with closed loop scanners using a liquid cell for imaging of supported lipid bilayers. All experiments were carried out at 20 °C. Tapping mode imaging was performed using aluminium coated cantilevers (PPP-NCSTR, Apex Probes, UK) and all images were obtained in PBS buffer. Contact mode images were obtained using contact mode aluminium coated silicon cantilevers (PPP-CONTR, Apex Probes, UK) having nominal spring constants of between 0.02 and 0.77 N m^-1^ and typical tip radius of curvatures of less than 7 nm.

### Amyloid beta preparation

Beta-amyloid (1-42), human (Anaspec) was reconstituted with 1.0% NH4OH and aliquoted before freezing. Before use the aliquot was defrosted and made up to the desired concentration with PBS buffer. This solution was then vortexed for 2 minutes before being added to the supported lipid bilayer. For all experiments 200 μl of 8 μM amyloid beta solution was added to the supported bilayer and left for 2 hr. This was removed by gentle pipetting and the bilayer was rinsed gently three times in PBS buffer to remove any free floating amyloid beta.

### Preparation of supported lipid bilayers

2-Dipalmitoyl-*sn*-glycero-3-phosphocholine (DOPC) and 1, 2-dioleoyl-*sn*-glycero-3-phosphocholine (DPPC) (Avanti Polar Lipids) were dissolved in chloroform in a 3 to 1 molar ratio and dried under nitrogen. These lipids mixtures were then re-suspended in ultrapure water to a final concentration of 1 mg/ml and intensively vortexed at room temperature. For brain derived lipids the lipids were dried down to form a film and then resuspended in ultrapure water. Uni-lamellar vesicles were formed via ten cycles of freeze-thawing followed by extrusion ten times) through a polycarbonate membrane with pores sizes of 100 nm using a mini-extruder (Avanti Polar Lipids) kept 50 °C on a hotplate. 200 μl of solution containing the extruded vesicles was pipetted onto freshly cleaved muscovite mica and this was then placed on a thin metal disk that itself was placed on a hotplate. The hotplate was heated up to 50 °C and the sample was annealed for 15 minutes. The sample was then loaded into the AFM liquid cell ready for imaging.

### Breakthrough force measurements

Indentation measurements were made using contact mode aluminium coated silicon cantilevers (PPP-CONTR, Apex Probes, UK) having nominal spring constants of between 0.02 and 0.77 N m^-1^ and typical tip radius of curvatures of less than 7 nm. The spring constants were calibrated using the equipartition theorem (Thermal K). Actual measured spring constant values were between 0.05 and 0.18 Nm^-1^. Force volume indentation grids of 256 points (16×16) were taken at no less than three different areas across the bilayer. Indentation rate was 1μm/s which was the optimum rate for measuring breakthrough events. Force distance curves were analyzed using a Scanning Probe Image Processor (Image Metrology, Lyngby, Denmark) and in house developed batch analysis code based on the methodology proposed by Li et al^32^.

### Statistical analysis

Data are presented as means ± standard error of the mean (SEM). Analyses were performed using GraphPad Prism 6.0 software (GraphPad, San Diego, CA). Comparisons between groups were performed by the Mann-Whitney test. Statistical significance was set at P < 0.05. The heatmaps were created with the R software (www.r-project.org) with R package, Marray. Ward’s algorithm, modified by Murtagh and Legendre, was used as the clustering method.

## Acknoweldgements

The authors would like to thank the University of Illinois at Chicago, Department of Chemistry and College of Liberal Arts and Science for support of this work. Funding was also received from: the National Science Foundation (Award #1655494 to T.J.P. and S.M.C.) and the UIC DFI Fellowship to M.R.P.; Cancer Resarch UK/RCUK (Award C56829/A22053 to D.F., K.R. and E.St.J.S.); BBSRC-DTP studentship (A.A.S.), Centre for Misfolding Diseases (A.A.S, S.P, M.V., J.R.K.); Agence Nationale de la Recherche (ANR-18-CE14-0039-01 to N.C). We thank Thu T.A. Nguyen for technical assistance.

