## Supplementary Info for "Lipid membranes from naked mole-rat brain lipids are cholesterol-rich, highly phase-separated, and sensitive to amyloid-induced damage"

**Supplementary materials**

### Production of naked mole-rat amyloid beta

The single point mutation (His13Arg) was introduced to the pET-Sac vector containing a gene encoding for wild type, human Aβ(M1–42) using a QuikChange Site Directed Mutagenesis kit (Agilent UK Ltd., Cheshire, UK) and following the manufacturer's protocol. The primers were: Forward-5'-GGTTACGAAGTTCGCCACCAGAAGCTGG-3' and Reverse - 5'-CCAGCTTCTGGTGGCGAACTTCGTAACC-3'. Plasmid sequences were confirmed by DNA sequencing (Dept. of Biochemistry, University of Cambridge, UK). The recombinant Naked Mole-Rat Aβ(M1-42) peptide (MDAEFRHDSGY EVRHQKLVFFAEDVGSNKGA IIGLMVGGVV IA), here called NMR amyloid beta, was expressed in the E. coli BL21 Gold (DE3) strain (Stratagene, CA, U.S.A.) and purified as described previously with slight modifications[1]. Briefly, the purification procedure involved sonication of E. coli cells, dissolution of inclusion bodies in 8 M urea, and ion exchange in batch mode on diethylaminoethyl cellulose resin followed by lyophylisation. The lyophilised fractions were further purified using Superdex 75 HR 26/60 column (GE Healthcare, Buckinghamshire, U.K.) and eluates were analysed by SDS-PAGE using 4-12% Bis-Tris NuPAGE gels and MES buffer (Thermofisher, Paisley, UK) for the presence of the desired protein product. The fractions containing the recombinant protein were combined, frozen using liquid nitrogen, and lyophilised again. The mass of the NMR amyloid beta was confirmed by mass spectrometry (Department of Chemistry, University of Cambridge, UK) (calculated mass: 4664.3 Da; observed mass: 4663.4 ± 0.8 Da).

### Thioflavin T (ThT) Kinetic Assay

A sample of lyophilised naked mole-rat amyloid beta was dissolved in 1 ml GdnHCl (6 M, pH 8) and incubated on ice (1.5-2 hr). SEC was performed on the NMR Aβ42 using a Superdex 75 10/300 GL column (GE Healthcare, Amersham, UK). The buffer used for elution was a pH 8 sodium phosphate buffer (25mM $Na_2PO_4$, 0.2mM EDTA, pH 8) (flow rate 0.7 ml/min). Solutions containing monomer were collected and monomer concentrations were determined by the UV absorbance of the solution ($\varepsilon_{280\ nm}$ =1490 $M^{-1}cm^{-1}$). The monomer obtained in this way was diluted with buffer to the desired concentration and supplemented with 20 µM Thioflavin T (ThT) from a 1 mM stock. All samples were prepared in low binding eppendorf tubes on ice using careful pipetting to avoid introduction of air bubbles. Each sample was then pipetted into multiple wells of a 96-well half-area, low-binding, clear bottom and PEG coated plate (Corning 3881), to give 80 µL per well. Kinetics assays were initiated by placing the 96-well plate at 37 ºC under quiescent conditions in a plate reader (Fluostar Omega, BMGLabtech, Offenburg, Germany). The ThT fluorescence was measured through the bottom of the plate with a 440 nm excitation filter and a 480 nm emission filter. The ThT fluorescence was followed for three repeats of each sample.

### Transmission Electron Microscopy

5µl of NMR and human Aβ42 aggregate solutions were applied to carbon-coated copper grids, stained with 2% (w/v) uranyl acetate and washed twice with 5 µl MilliQ water. The grids were then imaged with a Thermo Scientific Talos F200X G2 200 kV FEG Scanning Transmission Electron Microscope (Department of Chemistry, University of Cambridge).

**Supplementary Table 1: percentage of phosphatidylcholine from mouse and naked mole rat brains.** Values are expressed as mean ± SEM; Statistical analysis was performed using Mann-Whitney test.

| | % of total phosphatidylcholine | | |
|---|---|---|---|
| Lipid number | Naked mole rat | Mouse | P value |
| PC 28:0 | 0.87 ± 0.12 | 0.021 ± 0.003 | 0.0012 |
| PC 30:0 | 8.80 ± 0.78 | 0.91 ± 0.10 | 0.0012 |
| PC 30:1 | 2.70 ± 1.04 | 0.59 ± 0.26 | 0.2343 |
| PC 32:0 | 22.04 ± 0.55 | 32.98 ± 0.58 | 0.0012 |
| PC 32:1 | 5.00 ± 0.86 | 1.08 ± 0.24 | 0.0012 |
| PC 32:2 | 0.14 ± 0.007 | 0.008 ± 0.0007 | 0.0012 |
| PC 34:0 | 2.90 ± 0.09 | 3.41 ± 0.10 | 0.0082 |
| PC 34:1 | 18.45 ± 0.61 | 17.67 ± 0.44 | 0.4452 |
| PC 34:2 | 1.50 ± 0.10 | 0.66 ± 0.02 | 0.0012 |
| PC 34:3 | 0.15 ± 0.01 | 0.02 ± 0.001 | 0.0012 |
| PC 36:1 | 4.19 ± 0.09 | 7.68 ± 0.08 | 0.0012 |
| PC 36:2 | 2.57 ± 0.19 | 2.98 ± 0.17 | 0.1014 |
| PC 36:3 | 1.51 ± 0.06 | 1.20 ± 0.03 | 0.0012 |
| PC 36:4 | 6.47 ± 0.12 | 5.69 ± 0.08 | 0.0012 |
| PC 38:2 | 0.21 ± 0.02 | 0.50 ± 0.01 | 0.0012 |
| PC 38:3 | 1.21 ± 0.02 | 1.07 ± 0.01 | 0.0012 |
| PC 38:4 | 7.83 ± 0.35 | 6.93 ± 0.25 | 0.1375 |
| PC 38:5 | 3.70 ± 0.10 | 2.43 ± 0.33 | 0.0012 |
| PC 38:6 | 3.98 ± 0.05 | 5.68 ± 0.20 | 0.0012 |
| PC 40:3 | 0.097 ± 0.002 | 0.08 ± 0.002 | 0.0140 |
| PC 40:6 | 5.64 ± 0.07 | 8.35 ± 0.26 | 0.0012 |

**Supplementary Table 2: percentage of phosphatidylethanolamine from mouse and naked mole rat brains.** Values are expressed as mean ± SEM; Statistical analysis was performed using Mann-Whitney test.

| | % of total phosphatidylethanolamine | | |
|---|---|---|---|
| Lipid number | Naked mole rat | Mouse | P value |
| PE 32:0 | 0.50 ± 0.01 | 0.39 ± 0.03 | 0.0140 |
| PE 32:1 | 0.56 ± 0.07 | 0.18 ± 0.02 | 0.0012 |
| PE 32:2 | 0.040 ± 0.004 | 0.007 ± 0.001 | 0.0012 |
| PE 34:0 | 8.88 ± 1.12 | 8.39 ± 1.02 | 0.2343 |
| PE 34:1 | 1.04 ± 0.15 | 0.57 ± 0.06 | 0.0221 |
| PE 36:1 | 8.46 ± 0.64 | 14.50 ± 0.40 | 0.0012 |
| PE 36:2 | 4.71 ± 0.76 | 7.32 ± 0.88 | 0.0350 |
| PE 36:3 | 1.77 ± 0.11 | 1.31 ± 0.16 | 0.1014 |
| PE 36:4 | 1.79 ± 0.09 | 1.42 ± 0.07 | 0.0140 |
| PE 38:2 | 1.83 ± 0.17 | 1.90 ± 0.10 | 0.9452 |
| PE 38:3 | 13.78 ± 2.55 | 8.26 ± 1.77 | 0.1014 |
| PE 38:4 | 16.55 ± 0.22 | 11.46 ± 0.22 | 0.0012 |
| PE 38:5 | 3.81 ± 0.43 | 5.25 ± 0.54 | 0.1014 |
| PE 38:6 | 4.40 ± 0.24 | 4.77 ± 0.53 | 0.9452 |
| PE 40:3 | 3.57 ± 0.65 | 1.96 ± 0.38 | 0.1014 |
| PE 40:5 | 12.14 ± 1.35 | 13.0& ± 2.26 | 0.7308 |
| PE 40:6 | 14.60 ± 1.66 | 17.85 ± 1.80 | 0.1014 |
| PE 40:7 | 1.55 ± 0.30 | 1.42 ± 0.26 | 0.2343 |

**Supplementary Table 3: percentage of phosphatidylinositol from mouse and naked mole rat brains.** Values are expressed as mean ± SEM; Statistical analysis was performed using Mann-Whitney test.

| | % of total phosphatidylinositol | | |
|---|---|---|---|
| Lipid number | Naked mole rat | Mouse | P value |
| PI 32:1 | 0.19 ± 0.02 | 0.147 ± 0.008 | 0.1014 |
| PI 34:1 | 1.40 ± 0.19 | 2.08 ± 0.06 | 0.0350 |
| PI 34:2 | 0.39 ± 0.07 | 0.38 ± 0.02 | 0.4452 |
| PI 36:0 | 0.24 ± 0.02 | 0.66 ± 0.04 | 0.0012 |
| PI 36:1 | 0.81 ± 0.09 | 2.10 ± 0.07 | 0.0012 |
| PI 36:2 | 0.67 ± 0.08 | 0.90 ± 0.02 | 0.1807 |
| PI 36:3 | 2.36 ± 0.11 | 2.58 ± 0.20 | 0.4452 |
| PI 38:1 | 0.10 ± 0.01 | 0.285 ± 0.008 | 0.0012 |
| PI 38:2 | 1.02 ± 0.04 | 1.04 ± 0.05 | 0.7308 |
| PI 38:3 | 18.49 ± 1.08 | 17.34 ± 1.04 | 0.1375 |
| PI 38:4 | 55.34 ± 0.38 | 52.74 ± 0.27 | 0.0012 |
| PI 38:5 | 16.58 ± 0.87 | 14.51 ± 0.99 | 0.2343 |
| PI 40:2 | 0.026 ± 0.001 | 0.032 ± 0.008 | 0.9452 |
| PI 40:3 | 0.118 ± 0.003 | 0.159 ± 0.007 | 0.0012 |
| PI 40:4 | 0.377 ± 0.009 | 0.51 ± 0.01 | 0.0012 |
| PI 40:5 | 0.73 ± 0.05 | 1.39 ± 0.05 | 0.0012 |
| PI 40:6 | 1.16 ± 0.15 | 3.13 ± 0.15 | 0.0012 |

**Supplementary Table 4: percentage of phosphatidylserine from mouse and naked mole rat brains.** Values are expressed as mean ± SEM; Statistical analysis was performed using Mann-Whitney test.

| | % of total phosphatidylserine | | |
|---|---|---|---|
| Lipid number | Naked mole rat | Mouse | P value |
| PS 32:0 | 0.027 ± 0.001 | 0.011 ± 0.001 | 0.0023 |
| PS 34:0 | 0.57 ± 0.02 | 0.48 ± 0.01 | 0.0047 |
| PS 34:1 | 2.37 ± 0.12 | 1.58 ± 0.02 | 0.0012 |
| PS 34:2 | 0.19 ± 0.01 | 0.091 ± 0.002 | 0.0012 |
| PS 36:0 | 2.03 ± 0.16 | 3.27 ± 0.08 | 0.0012 |
| PS 36:1 | 6.66 ± 0.36 | 12.22 ± 0.37 | 0.0012 |
| PS 36:2 | 3.32 ± 0.14 | 4.88 ± 0.16 | 0.0012 |
| PS 36:3 | 0.345 ± 0.007 | 0.188 ± 0.003 | 0.0012 |
| PS 38:2 | 0.74 ± 0.03 | 0.86 ± 0.04 | 0.0350 |
| PS 38:3 | 2.57 ± 0.10 | 1.84 ± 0.06 | 0.0023 |
| PS 38:4 | 5.29 ± 0.07 | 3.71 ± 0.13 | 0.0012 |
| PS 38:6 | 0.38 ± 0.04 | 0.61 ± 0.06 | 0.0221 |
| PS 40:5 | 23.33 ± 0.43 | 18.02 ± 0.21 | 0.0012 |
| PS 40:6 | 51.91 ± 0.43 | 52.01 ± 0.32 | 0.5338 |
| PS 42:6 | 0.263 ± 0.009 | 0.23 ± 0.01 | 0.1807 |

**Supplementary Table 5: percentage of sphingomyelin from mouse and naked mole rat brains.** Values are expressed as mean ± SEM; Statistical analysis was performed using Mann-Whitney test.

| | % of total sphingomyelin | | |
|---|---|---|---|
| Lipid number | Naked mole rat | Mouse | P value |
| SM 18:1/14:0 | 1.95 ± 0.07 | 0.113 ± 0.008 | 0.0012 |
| SM 18:1/16:0 | 20.05 ± 0.95 | 4.16 ± 0.18 | 0.0012 |
| SM 18:1/16:1 | 0.46 ± 0.05 | 0.131 ± 0.009 | 0.0012 |
| SM 18:1/18:0 | 58.86 ± 0.68 | 68.62 ± 1.05 | 0.0012 |
| SM 18:1/18:1 | 5.46 ± 0.54 | 5.79 ± 0.39 | 0.9452 |
| SM 18:1/20:0 | 1.61 ± 0.14 | 2.53 ± 0.31 | 0.0734 |
| SM 18:1/20:1 | 0.150 ± 0.009 | 0.35 ± 0.02 | 0.0012 |
| SM 18:1/22:0 | 1.08 ± 0.18 | 1.54 ± 0.16 | 0.1014 |
| SM 18:1/22:1 | 0.56 ± 0.05 | 0.83 ± 0.13 | 0.2949 |
| SM 18:1/24:0 | 2.83 ± 0.33 | 3.03 ± 0.11 | 0.7308 |
| SM 18:1/24:1 | 6.98 ± 0.77 | 12.90 ± 0.52 | 0.0012 |

**Supplementary Table 6: percentage of ceramide from mouse and naked mole rat brains.** Values are expressed as mean ± SEM; Statistical analysis was performed using Mann-Whitney test.

| | % of total ceramide | | |
|---|---|---|---|
| Lipid number | Naked mole rat | Mouse | P value |
| Cer d18:1/16:0 | 12.39 ± 0.66 | 1.99 ± 0.06 | 0.0012 |
| Cer d18:1/16:1 | 0.087 ± 0.003 | 0.018 ± 0.001 | 0.0012 |
| Cer d18:1/18:0 | 77.92 ± 0.38 | 85.18 ± 1.70 | 0.0012 |
| Cer d18:1/18:1 | 1.15 ± 0.07 | 0.91 ± 0.09 | 0.1375 |
| Cer d18:1/20:0 | 0.76 ± 0.03 | 3.40 ± 0.08 | 0.0012 |
| Cer d18:1/22:0 | 1.11 ± 0.06 | 1.35 ± 0.10 | 0.1014 |
| Cer d18:1/24:0 | 3.06 ± 0.44 | 3.12 ± 0.71 | >0.9999 |
| Cer d18:1/24:1 | 3.45 ± 0.47 | 3.97 ± 0.99 | 0.8357 |
| Cer d18:1/26:0 | 0.033 ± 0.008 | 0.023 ± 0.008 | 0.2343 |
| Cer d18:1/26:1 | 0.038 ± 0.007 | 0.022 ± 0.006 | 0.1014 |

**Supplementary Table 7: Relative concentration of fatty acid from mouse and naked mole rat brains.** Values are expressed as mean ± SEM; Statistical analysis was performed using Mann-Whitney test.

| | | Relative concentration per g of tissue | | |
|---|---|---|---|---|
| Common name | Lipid number | Mouse | Naked mole rat | P value |
| Myristic acid | C14:0 | 0.84 ± 0.08 | 2.53 ± 0.19 | 0.0025 |
| Palmitic acid | C16:0 | 0.73 ± 0.02 | 1.14 ± 0.05 | 0.0012 |
| Stearic acid | C18:0 | 670.90 ± 69.27 | 526.00 ± 33.00 | 0.1375 |
| Hypogeic acid | C16:1 n-9 | 3.03 ± 0.50 | 17.90 ± 2.14 | 0.0012 |
| Palmitoleic acid | C16:1 n-7 | 8.62 ± 1.25 | 11.00 ± 1.13 | 0.2774 |
| Elaidic acid | C18:1 n-9 | 540.60 ± 59.06 | 425.70 ± 21.36 | 0.1807 |
| Vaccenic acid | C18:1 n-7 | 117.50 ± 13.06 | 110.20 ± 5.10 | 0.6282 |
| Gondoic acid | C20:1 n-9 | 75.29 ± 11.01 | 35.49 ± 6.50 | 0.0140 |
| Linoleic acid | C18:2 n-6 | 20.48 ± 2.86 | 22.66 ± 2.13 | 0.8357 |
| α-Linolenic acid | C18:3 n-3 | 0.87 ± 0.14 | 0.87 ± 0.14 | 0.9452 |
| Eicosadienoic acid | C20:2 n-6 | 8.77 ± 1.09 | 9.39 ± 1.03 | 0.7308 |
| Dihomo-γ-linolenic acid | C20:3 n-6 | 19.39 ± 2.25 | 21.26 ± 1.79 | 0.5589 |
| Arachidonic acid | C20:4 n-6 | 353.40 ± 42.06 | 360.10 ± 29.88 | 0.8357 |
| Eicosapentaenoic acid | C20:5 n-3 | 1.11 ± 0.15 | 5.40 ± 0.89 | 0.0012 |
| Docosatrienoic acid | C22:3 n-3 | 7.81 ± 1.17 | 24.20 ± 3.36 | 0.0023 |
| Adrenic acid | C22:4 n-6 | 89.30 ± 11.63 | 93.05 ± 10.27 | 0.9999 |
| Docosapentaenoic acid | C22:5 n-3 | 11.77 ± 2.09 | 38.43 ± 3.11 | 0.0012 |
| Docosahexaenoic acid | C22:6 n-3 | 660.00 ± 88.74 | 382.30 ± 1.67 | 0.0221 |

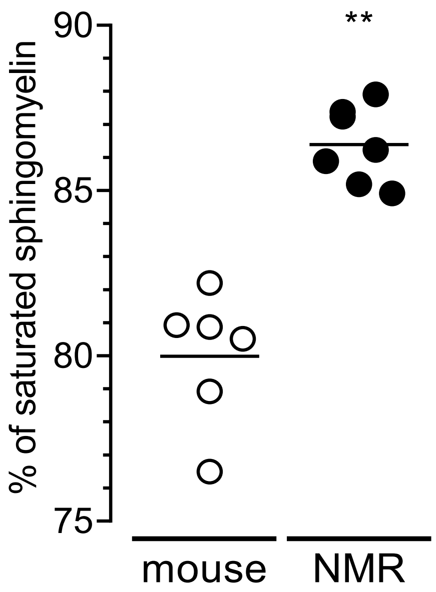


**Supplementary Figure 1: Percentage of saturated sphingomyelin. Lipids were extracted from the brains of 7 naked mole-rats and 6 mice (2 independent experiments). % of saturated sphingomyelin in lipid extract from mouse (white circle) or naked mole-rat (black circle) brains. Data are represented as scatter dot plot with the mean. Statistical analysis was performed using the Mann-Whitney test. **p<0.01, significantly different from mouse group.**

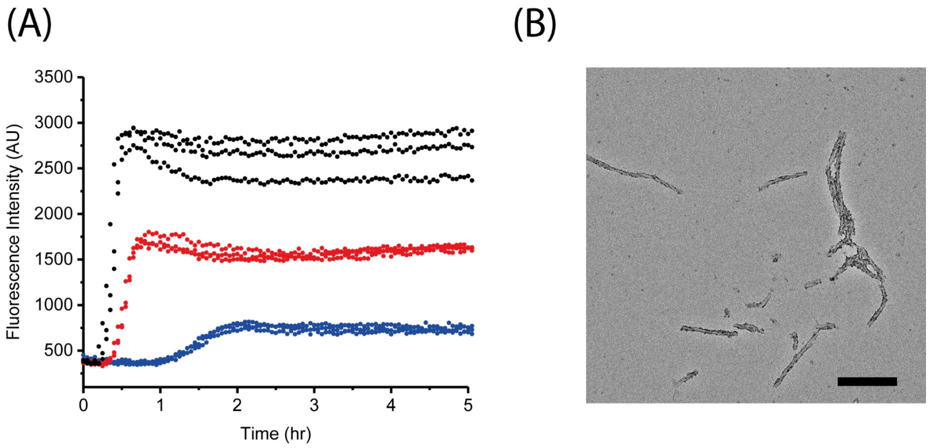


**Supplementary Figure 2: A) ThT kinetics traces for 1 µM (blue), 3 µM (red) and 5 µM (black) naked mole-rat amyloid beta shown in triplicate (pH 8, 37°C, quiescent conditions). Representative of two independent experiments. B) TEM analysis of endpoint fibrils formed from kinetics assay (scale bar = 200 nm). Results indicative of a single batch of naked mole-rat amyloid beta.**

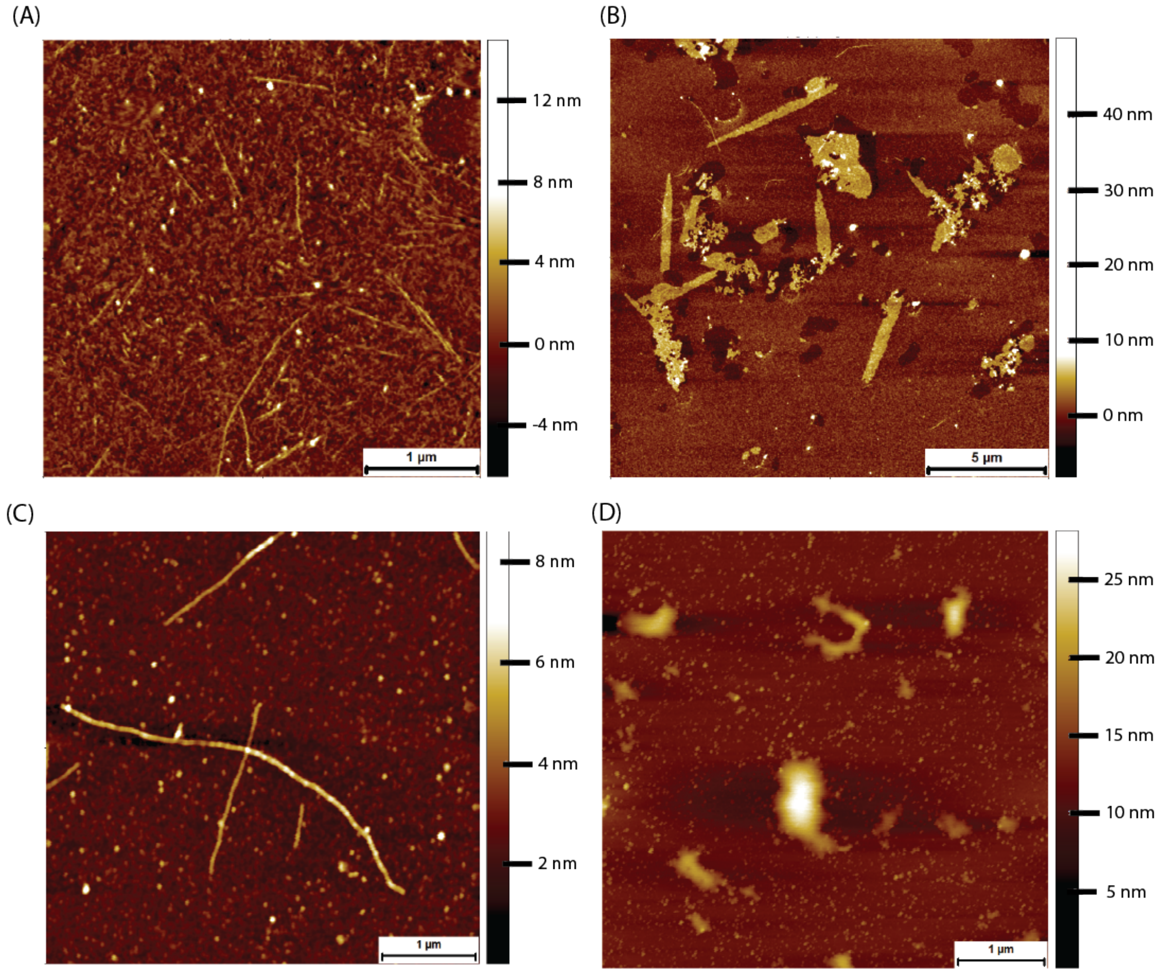


**Supplementary Figure 3 – Amyloid beta peptide adsorbed (air dried) onto mica at a concentration of 8 µM. A) Synthetic human amyloid beta readily forms fibres. B) Higher order structures of the human amyloid beta. These are well organised, and the constituent fibres are resolvable. C) Recombinantly expressed naked mole-rat amyloid beta also forms fibres. D) Less well-defined**

**higher order structures are formed by naked mole-rat amyloid beta.**

(A)

(B)

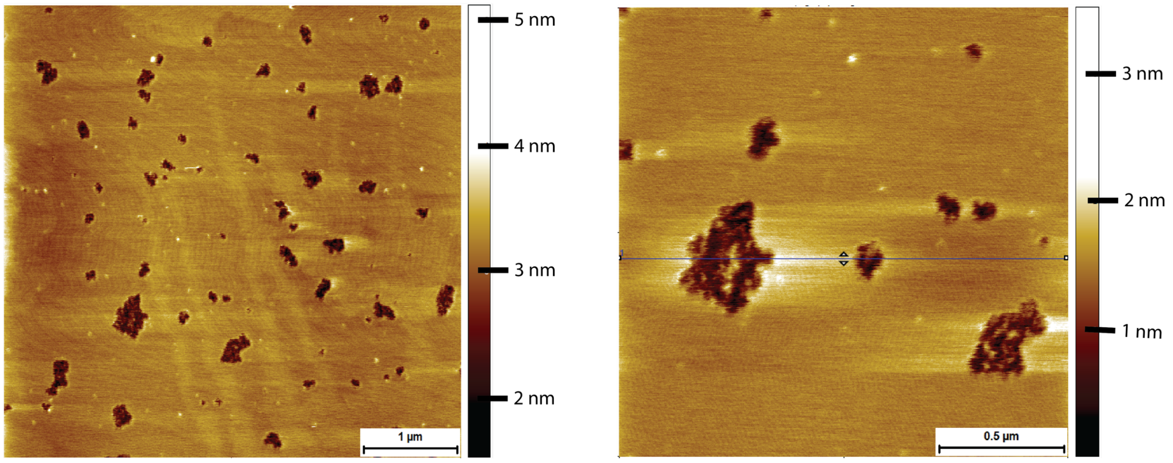


**Supplementary Figure 4 – AFM tapping mode images (in PBS) of mouse brain tissue derived supported lipid bilayers exposed to 8 μM of naked mole-rat amyloid beta for 2 hr. A) Supported bilayer made from mouse brain derived lipids exhibiting naked mole-rat amyloid beta induced holes. B) High resolution tapping mode image of naked mole-rat amyloid beta induced holes in mouse brain lipid derived bilayers. Holes do not penetrate bilayer.**

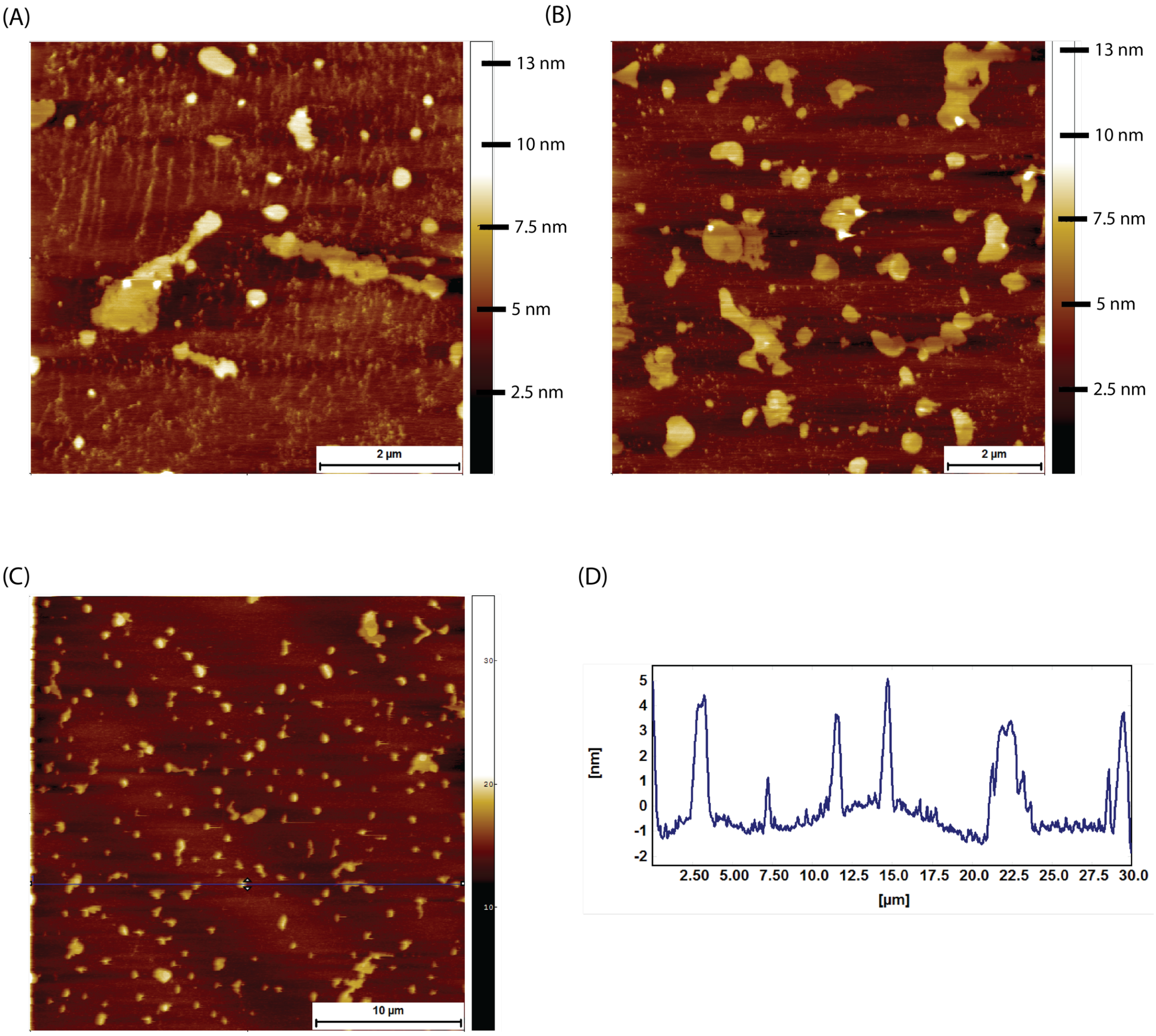


**Supplementary Figure 5 – AFM tapping mode images (in PBS) of naked mole-rat brain derived lipid bilayers exposed to 8 µM of naked mole-rat amyloid beta for 2 hr. (A and B) The bilayer is reduced to fragments of membrane, the height of fragments consistent with bilayer thickness. (C) Large scale topographic image showing the scale of the damage. (D) A height profile corresponding to the blue horizontal line in (C), confirming the height of fragments to be of bilayer thickness of 5-6 nm.**

1 Hellstrand, E., Boland, B., Walsh, D. M. & Linse, S. Amyloid beta-protein aggregation produces highly reproducible kinetic data and occurs by a two-phase process. *ACS Chem Neurosci* **1**, 13-18, doi:10.1021/cn900015v (2010).
